# Identifying high-priority proteins across the human diseasome using semantic similarity

**DOI:** 10.1101/309203

**Authors:** Edward Lau, Vidya Venkatraman, Cody T Thomas, Jennifer E Van Eyk, Maggie PY Lam

**Author notes:** Correspondence Maggie Pui Yu Lam University of Colorado Denver - Anschutz Medical Campus Mail Stop B139, Research Complex 2 12700 E. 19th Avenue Aurora, CO 80045.

## Abstract

Knowledge of “popular proteins” has been a focus of multiple Human Proteome Organization (HUPO) initiatives and can guide the development of proteomics assays targeting important disease pathways. We report here an updated method to identify prioritized protein lists from the research literature, and apply it to catalog lists of important proteins across multiple cell types, sub-anatomical regions, and disease phenotypes of interest. We provide a systematic collection of popular proteins across 10,129 human diseases as defined by the Disease Ontology, 10,642 disease phenotypes defined by Human Phenotype Ontology, and 2,370 cellular pathways defined by Pathway Ontology. This strategy allows instant retrieval of popular proteins across the human “diseasome”, and further allows reverse queries from protein to disease, enabling functional analysis of experimental protein lists using bibliometric annotations.

## Introduction

The corpus of scientific literature presents a network of structured associations among genes/proteins, diseases, and researchers [Fortunato et al., 2018]. Identifying the prioritized proteins within a biomedical topic, whether it be a disease, cell type, or organ, can yield insights into the research trends and important pathways within a biomedical area of interest. Recently it has been recognized that understanding protein-disease relationships provides a valuable tool to guide proteomics researchers to prioritize their efforts on developing proteomics assays to examine proteins that are of interest to a board number of researchers. We previously demonstrated a method and workflow to objectively interrogate the most popular proteins in a number of research areas and organ systems [Lam et al., 2016a,b]. Our workflow is based on measurements of semantic similarity between genes/proteins and literature publications using publicly available resources provided by the search function on PubMed and a gene to PMID reference dataset available on NCBI. By combining these online resources, we were able to determine the most published proteins/genes in 6 organ systems and beyond. The Biology/Disease Human Proteome Project (B/D-HPP) initiatives within the Human Proteome Organization (HUPO) have widely adopted this approach to discover critical proteins in the heart [Lam et al., 2015], liver [Mora et al., 2017], brain [Lam et al., 2016b], and eye [Semba et al., 2015], and the results of which are leveraged to identify research trends and prioritize bioassay development.

Here we have expanded on the original normalized co-publication distance (NCD) approach. The updated approach incorporates additional data sources and further introduces a regularized copublication distance (RCD) metric which takes into account the immediacy and influence of individual publications to limit outsized contributions of single publications to influence popularity scores. We show that this approach outperforms the published NCD scores. We also compare this improved approach with parallel efforts to identify prioritized gene lists using literature data. We demonstrate the utility of the approach to identify popular proteins across diseases and disease phenotypes, including inflammation, fibrosis, metabolic syndrome, protein misfolding, and cell death. We then extended this approach to identify significant protein lists from a compilation of known human diseases sometimes collectively referred to as the human “diseasome” or disease network [Hoehndorf et al., 2015, Zhou et al., 2014], by querying a vast collection of over 23,000 biomedical terms compiled in standardized vocabularies of human disease processes including 10,129 diseases, and 10,642 phenotypes, and 2,370 pathways. We show that disease search terms are associated with specific prioritized protein lists whereas similar diseases are associated with similar prioritized proteins. Finally, we have implemented a reverse protein search strategy over the precompiled terms, which associates an input list of genes/proteins with the diseases and disease phenotypes in which they are intensively investigated.

## Materials and Methods

### Calculation of semantic distance between protein and topics

The popular protein strategy performs large-scale bibliometric analysis from the research papers curated on PubMed. Research topics are used to query PubMed via the NCBI EUtils Application Programming Interfaces [Agarwala et al., 2018] to retrieve associated articles, and an annotation table that houses known PubMed ID (PMID)-Gene associations. To measure the semantic distance between a gene/protein with a topic of interest in the literature, we previously devised a semantic similarity metric NCD, defined as:

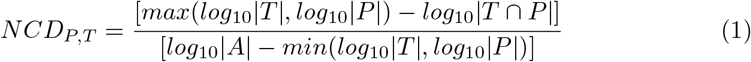

where *T* denotes the set of articles associated with any protein in the set of articles contained in the annotation table and that are retrieved from the PubMed query; *P* is the set of articles associated with a particular protein in the annotation table; *A* is the set of all articles associated with any proteins in any topics in the annotation, i.e., all PubMed IDs in the annotation table; such *T* ⊆ *A* and *P* ⊆ *A*.

Here we devised a regularized variant of NCD by introducing weighted adjustments to each article’s contribution by immediacy and influence metrics. In the unadjusted NCD each associated article *a_i_* carries an equal weight of 1. The weight is adjusted in the regularized co-publication distance (RCD) such that each annotated article in the association table i carries a weight of *w_i_*:

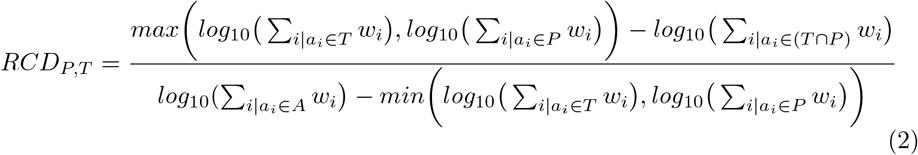

To model the influence of a publication *a_i_*, we applied the logistic transformation of the base 10 logarithm of the number of citations of a publication plus one *n_i_*, where the scale *a*, shape *b* and steepness *c* are 1, 6 and 2, respectively.

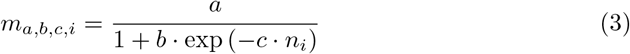

To model the immediacy of a paper, we use a weibull transformation of distance in decades since the publication date of the paper to the present *y_i_*, with the shape parameters λ and *k* heuristically set to 1 and 1.25.

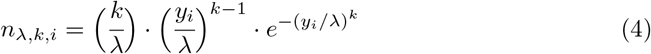

The final weight of the paper is calculated as the sum of the associated publication counts plus each associated publication’s immediacy and influence.

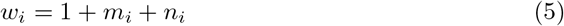

### Determination of popular protein lists

With the above method, we retrieved popular protein lists with search terms as described in the results section. Protein-PMID associations are retrieved on 2018-04-22 from NCBI Gene2Pubmed (NCBI Resource Coordinators 2018) curations, and data downloaded from Pubtator [Wei et al., 2013], the latter of which contains text-mined relationships between biomedical concepts and entities. A union of the two sets of curation was used.

We performed PubMed queries on 27,257 defined topics retrieved from publicly available vocabularies, including 11,093 disease definitions from Disease Ontology (DOID) [Kibbe et al., 2015] (version 2018-03-02; retrieved 2018-03-15), 13,537 phenotypic descriptions from Human Phenotype Ontology (HPO) [Kohler et al., 2017] (version 2018-03-08; retrieved 2018-03-15), and 2,627 biochemical and signaling pathways from Pathway Ontology (PW) [Petri et al., 2014] (version 7-4-2; retrieved 2018-03-15). From the retrieved PubMed IDs from each query, protein-term associations are ranked according to normalized co-publication distance as previously described (popularity index). Moreover, the popularity index for all terms and proteins that are significantly associated with each individual topics (*P* < 0.05) have been uploaded to the PubPular web app and are made searchable.

### Web application and user interface

We provide a web app Pubpular (http://pubpular.org) that allows users to query the popular protein lists of custom topics. The PubPular web app automatically analyzes the occurrences of each protein being referenced to the retrieved papers using the Gene2PubMed [Agarwala et al., 2018] and Pubtator [Wei et al., 2013] resources, and performs calculation of RCD between a protein and the queried topic. We created an extended module to the PubPular web application named FABIAN (Functional Annotation by Bibliometric Analysis) which provides the functionality for gene lists to be uploaded and compared to results from curated terms. The module builds on the precompiled results from search terms on PubPularDB and uses parametric gene set enrichment analysis [Kim and Volsky, 2005] to discover terms for which the list of associated proteins (*P* ≤ 0.05) are significantly enriched or depleted with reference to the ranks of the uploaded gene list.

### Comparison of prioritized gene lists against curated standards

Curated “gold standard” protein lists were retrieved as follows: Proteins associated with three Gene Ontology (GO) [Consortium., 2017] terms for disease processes namely apoptosis (apoptotic process, GO:0006915; 762 genes), cell adhesion (GO:0007155; 800 genes), and DNA repair (463 genes; GO:0006281) were retrieved from European Bioinformatics Institute (EBI) QuickGO interface [Binns et al., 2009] and filtered to include only human Entrez Gene IDs that exist in the annotation source. Proteins associated with three complex disease terms namely hypertension (170 genes), obesity (202 genes), and schizophrenia (180 genes) were retrieved from Comparative Toxicogenomics Database (CTD) [Grondin et al., 2018] (downloaded on 2018-04-21). Comparisons with the prioritized gene/protein lists from GLAD4U [Jourquin et al., 2012] and PURPOSE [Yu et al., 2018] were perform by downloading query results after accessing the web tools and entering the exact search terms as shown and the protein lists were retrieved in entirety using their download functions. Precision for A/B classification is defined as *trueA*/(*trueA* + *trueB*), recall is defined as *trueA*/(*trueA* + *falseB*) and *F_β_* is defined as (1 + *β*^2^) · (*precision · recall*)/(*β · precision* + *recall*).

## Results

### Evaluation of prioritized gene lists based on regularized co-publication distance

We previously devised a metric NCD for the semantic similarity between a protein and a topic of interest. NCD normalizes the count of query-specific publications by the count of total publications on PubMed that are associated with the protein, so that a query will not be populated only by proteins that are broadly studied in many fields (e.g., p53 or APOE). RCD modifies NCD by modeling the immediacy and influence of an article to weight its contribution to the overall protein-term association (Figure 1).

**Figure 1.**
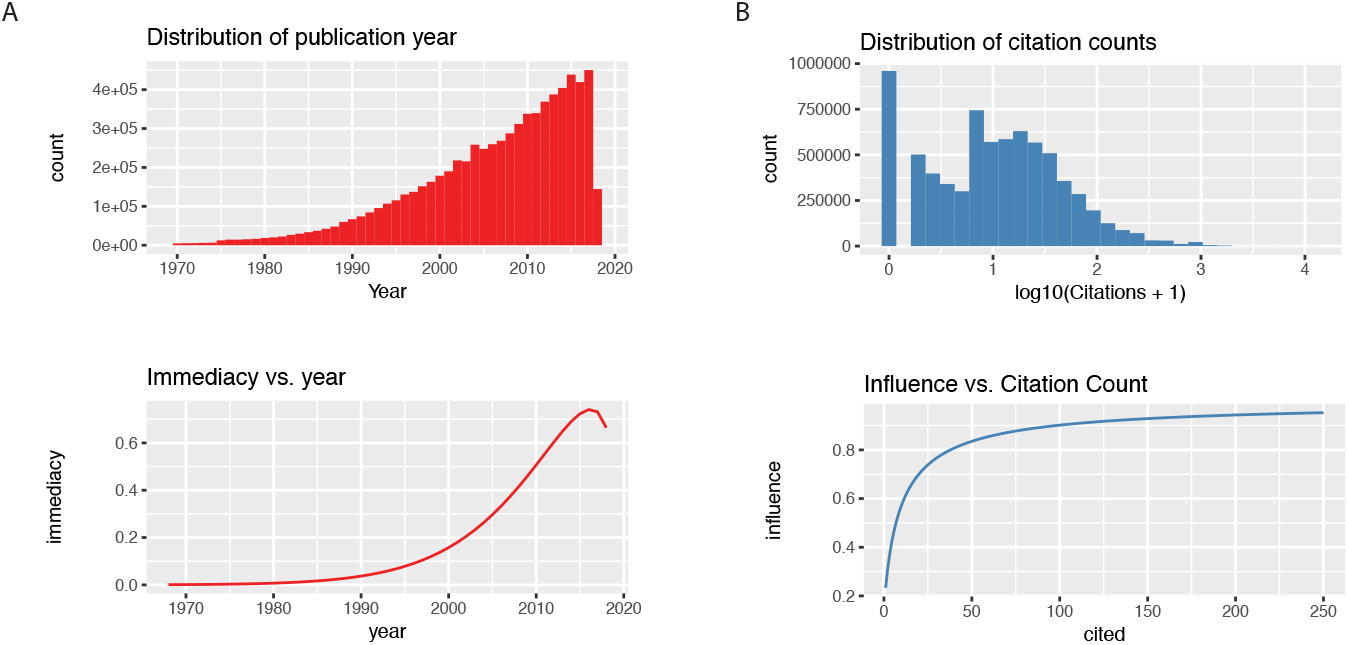
Modeling the immediacy and influence of protein-associated publications. **A** The immediacy of a publication is modeled using a weibull distribution such that recent publications within the past decade are given greater weights. **B** The influence of a publication is modeled using a logistic transformation of the log citation counts of the publication retrieved from Europe PubMed central.

The resulting metrics prioritizes top proteins in PubMed query search terms including searches for inherited and complex diseases (Figure 2). A lower RCD for a protein within a disease query suggests higher semantic similarity between the protein term and the disease term in the literature, and is overall correlated with greater number of publications for the proteins within that topic.

**Figure 2.**
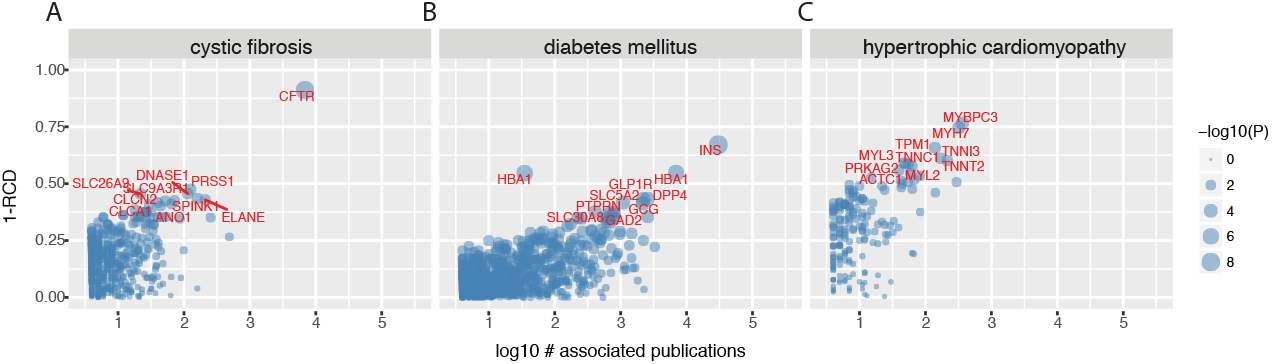
Prioritized protein lists in selected diseases. **A-C** Top prioritized protein lists in cystic fibrosis, diabetes mellitus, and hypertrophic cardiomyopathy are shown.

To compare the performance of RCD to retrieve relevant gene lists, we compared the results to a list of benchmark curated terms in public resources. These including curated biological process terms from Gene Ontology (cell adhesion, DNA repair, and apoptosis) as well as four complexe disease terms (hypertension, insulin resistance, obesity, and schizophrenia). Gene Ontology contains manually and automated curated relationship between terms and genes [Consortium., 2017], whereas the Comparative Toxicogenomics Database (CTD) [Grondin et al., 2018] collages annotations from Gene Ontology, Reactome, PubMed, and other sources. Both databases contain curated lists of good quality and are well utilized by researchers. We compared the performance to the NCD from the PubPular2 webapp NCD [Lam et al., 2016b] and to two other web tools namely GLAD4U [Jourquin et al., 2012] and PURPOSE [Yu et al., 2018]. The test terms were chosen to avoid biasing towards the present approach with specific terms as six of the seven terms were used as gold standards in the GLAD4U publication and the CTD database was a data source used to benchmark PURPOSE in its publication (Table 1).

**Table 1.** F-measure against benchmark curated gene lists.

| Term | Source | Protein | (Precision, recall, $F_1$, $F_2$) | | | |
| --- | --- | --- | --- | --- | --- | --- |
|  |  |  | GLAD4U | PURPOSE | PubPular NCD | RCD |
| Apoptosis | GO | 762 | 0.44, 0.40, 0.42, 0.41 | 0.08, 0.73, 0.16, 0.30 | 0.60, 0.30, 0.40, 0.33 | 0.49, 0.30, 0.37, 0.33 |
| Cell adhesion | GO | 800 | 0.51, 0.46, 0.49, 0.47 | 0.11, 0.56, 0.18, 0.30 | 0.69, 0.29, 0.41, 0.33 | 0.59, 0.34, 0.43, 0.37 |
| DNA repair | GO | 463 | 0.51, 0.58, 0.54, 0.56 | 0.12, 0.61, 0.19, 0.33 | 0.90, 0.39, 0.55, 0.44 | 0.81, 0.43, 0.56, 0.48 |
| Hypertension | CTD | 170 | 0.23, 0.38, 0.29, 0.34 | 0.05, 0.84, 0.09, 0.19 | 0.11, 0.31, 0.16, 0.23 | 0.20, 0.28, 0.23, 0.26 |
| Insulin resistance | CTD | 61 | 0.16, 0.62, 0.26, 0.40 | 0.03, 0.88, 0.05, 0.12 | 0.17, 0.30, 0.21, 0.25 | 0.15, 0.48, 0.23, 0.33 |
| Obesity | CTD | 202 | 0.23, 0.45, 0.30, 0.38 | 0.05, 0.70, 0.09, 0.18 | 0.14, 0.36, 0.20, 0.28 | 0.22, 0.29, 0.25, 0.27 |
| Schizophrenia | CTD | 180 | 0.16, 0.45, 0.24, 0.33 | 0.05, 0.59, 0.10, 0.19 | 0.34, 0.15, 0.22, 0.30 | 0.31, 0.20, 0.24, 0.20 |

We evaluate each prioritized protein list by its *F*-measure, which is a function of recall and precision of the approaches. The *F*_1_ score is equal to the harmonic mean of precision and recall to take into account the accuracy and specificity of prioritized protein lists, such that an indiscriminate prioritized protein list containing all known proteins will have high recall but low precision. The *F*_2_ score places twice the emphasis on recall over precision. We find that the compared methods differ in precision vs. recall across disease processes vs. complex disease terms. Overall the RCD-based prioritization improves on the F1 measure of four out of six terms against NCD. The precision and recall of the prioritized lists compares with similar approaches including GLAD4U and PURPOSE. We note that each strategy achieves different trade offs for recall vs. precision.

### Catalogs of popular proteins across cell types, and diseases

Using the devised prioritization method, we set out to identify prioritized proteins in several individual sub-anatomical regions and cell types. We previously demonstrated that queries of six major organ systems revealed specific affinity with different proteins. Here we asked whether the sub-anatomical regions and cell types can also be shown to be preferentially associated with different proteins. For the heart we queried the anatomical regions “left atrium”, “left ventricle”, “right atrium”, and “right ventricle” as well as the cell types “cardiomyocytes”, “smooth muscle cells”, “endothelial cells”, and “fibroblasts”, using the search terms “cardiac OR heart AND left AND atrium”, etc. For the lung, we queried the anatomical regions “alveolar sac”, “bronchiole”, “capillaries”, as well as the cell types “pneumocytes”, “smooth muscle”, “epithelial”, and “fibroblasts”. From the brain, we queried proteins associated with the anatomical regions “cerebellum”, “cerebrum”, and “brain stem”, and the cell types “neurons”, “astrocyte”, “glial cell”, and “oligodendrocyte”.

The analysis led to several general observations (Table 2). Firstly, we found that queries of different sub-anatomical regions were sufficiently specific and returned associations with region-specific proteins. For example, in the heart connexin-40 (GJA5) is preferentially associated with the atria but not ventricles, consistent with the known involvement of the protein in the pathogenesis of atrial fibrillation [van der Velden and Jongsma, 2002]. In the brain, ataxins (ATXN1/2), associated with progressive ataxias; are preferentially associated with the cerebellum but not the cerebrum. Cell types from each tissues were also associated with different lists of prioritized proteins. For instance, surfactant proteins are preferentially associated with pneumocytes, which form the alveolar linings, whereas fibroblast growth factors (FGFs) populate the prioritized list for lung fibroblasts. Notably, the fibroblasts and smooth muscle cells in the heart and in the lung are found to be associated with different sets of proteins, e.g., FGF23 and FGF21 for heart fibroblasts vs. FGF10 and FGF7 for lung fibroblasts, suggesting the prioritized protein lists may help shed light into the gene expression and properties of similar cell types found across multiple organs, such as fibroblasts and endothelial cells, that may be implicated in common disease processes, e.g., fibrosis and I endothelial disorders, that accompany diverse human diseases.

**Table 2.** Prioritized proteins across cell types and anatomical regions.

|  | Cell Types |  |  |  | Anatomical Regions |  |  |  |
| --- | --- | --- | --- | --- | --- | --- | --- | --- |
| Heart | Cardiomyocytes | Smooth muscle cell | Endothelial cell | Fibroblast | Left atrium | Left ventricle | Right atrium | Right ventricle |
|  | TTN | MYOCD | NOS3 | FGF23 | NPPB | NPPA | PKP2 | NPPA |
|  | NKX2-5 | TAGLN | PECAM1 | FGF21 | TP53INP2 | NPPB | NPPB | GJA5 |
|  | GATA4 | SRF | KDR | FGF2 | ADRB1 | GJA5 | TBX1 | ADRB1 |
|  | RYR2 | ACTA1 | VCAM1 | MYOCD | MYH7 | MYL4 | DSG2 | HCN4 |
|  | SCN5A | KCNJ8 | CDH5 | GATA4 | MYBPC3 | HCN4 | HAND2 | GJC1 |
| Lung | Pneumocytes | Smooth muscle cell | Epithelial cell | Fibroblasts | Alveolar sac | Bronchiole | Capillaries |  |
|  | SFTPC | ACTA1A | CFTR | CFHR1 | FOXQ1 | SCGB1A1 | PLSCR2 |  |
|  | SFTPA1 | BMPR2 | SFTPC | FGF10 | FOXJ1 | CYP2F1 | PLEK2 |  |
|  | SFTPB | EDN1 | CDH1 | FGF7 | FOXF1 | KCNRG | GPR182 |  |
|  | SFTPD | KCNA5 | SCGB1A1 | TGFB1 | ATP1A1 | SAA2-SAA4 | COL15A1 |  |
|  | NKX2-1 | KCNS3 | CXCL8 | FGFR1 | ABCA3 | SFTPB | PIANP |  |
| Brain | Neuron | Astrocyte | Glia | Oligodendrocyte | Cerebellum | Cerebrum | Brainstem |  |
|  | CA1 | GFAP | GFAP | OLIG2 | CBLN1 | CA1 | TH |  |
|  | CA3 | SLC1A2 | GDNF | MOG | CACNA1A | CA3 | OLIG2 |  |
|  | BDNF | AQP4 | OLIG2 | MBP | ATXN1 | PVALB | PROM1 |  |
|  | PVALB | SLC1A2 | AIF1 | CNP | ATXN2 | GRIA1 | GFAP |  |
|  | RBFOX3 | AIF1 | SLC1A2 | CSPG4 | GRID2 | BDNF | BDNF |  |

The majority of known human diseases can be grouped into subnetworks within a disease network sometimes called the “diseasome” an diseases can be grouped into clusters based on their shared disease phenotypes [Zhou et al., 2014]. To determine how the prioritized protein lists intersect with common disease processes that occur in complex human diseases including those that are the thematic focuses of HUPO B/D HPP initiatives, we queried six specific disease processes, namely fibrosis, cell death, inflammation, metabolic syndrome, oxidative stress, and protein misfolding (Table 3).

We find that the top five proteins in “fibrosis” are transforming growth factor beta 1 (TGFB1), followed by connective tissue growth factor (CTGF), actin alpha skeletal muscle (ACTA1), mothers against decapentaplegic homolog 3 (SMAD3), and mothers against decapentaplegic homolog 2 (SMAD2). Another molecular phenotype common in multiple diseases is “cell death”. The top 5 proteins in our popular protein search using the key cell death returned caspase-3 (CASP3), apoptosis regulator BAX (BAX), programmed cell death protein 1 (PDCD1), apoptosis regulator Bcl-2 (BCL2), and tumor necrosis factor ligand superfamily member 10 (TNFSF10). The query for inflammatory response returned common cytokines including interleukin-6 (IL6), C-reactive protein (CRP), tumor necrosis factor (TNF), interleukin-1 beta (IL1B), and interleukin-8 (CXCL8); the query for metabolic syndrome returned lipid metabolism proteins including adiponectin (ADPOQ), insulin (INS), fatty acid-binding protein (FABP4), and leptin (LEP); oxidative stress queries returned nuclear factor erythroid 2-related factor 2 (NRF2/NEF2L2), catalase (CAT), superoxide dismutases (SOD1/2), and kelch-like ECH-associated protein 1 (KEAP1). Finally protein misfolding returned tauopathy and neurodegenerative proteins as well as amyloidosis proteins including alternative prion protein (PRNP), alpha-synuclein (SNCA), huntingtin (HTT), transthyretin (TTR), and superoxide dismutase (SOD1).

**Table 3.**
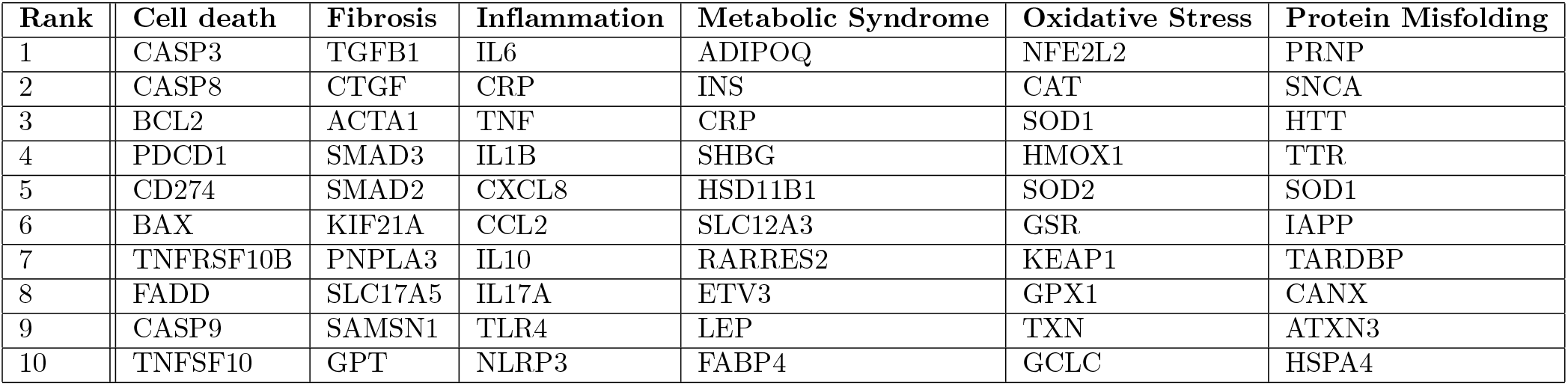
Top popular proteins for common disease phenotypes. Top 10 proteins represented by their gene name for each common disease characteristic are shown and ranked by RCD and their p-values.

### Protein-disease networks across the human diseasome

While the individual results on common disease processes are perhaps not surprising on their own, the prioritized gene lists could potentially be useful for identifying genes and pathways that are preferentially studied in particular disease processes such that reagent development efforts could be prioritized towards these topics (e.g., fibrosis in the heart). Building on this effort, we then systematically queried over 25,000 search terms in comprehensive vocabularies that describe the entirety of known human diseases (the “diseasome”). In total, we performed PubMed queries and calculated the list of prioritized proteins for 23,141 defined topics retrieved from publicly available vocabularies, including proteins for 10,129 disease definitions from Disease Ontology (DOID), 10,642 phenotypic descriptions from Human Phenotype Ontology (HPO), and 2,370 biochemical and signaling pathways from Pathway Ontology (PW). Among the vocabularies, 6,214 DOID terms are associated with no fewer than 50 distinct proteins, along with 5,342 in HPO and 1,479 in PW. Moreover, 7,897 search terms in DOID were associated with at least one significant (*P* ≤ 0.05) protein; along with 7,076 terms in HPO and 1,798 terms in PW (Figure 3.)

**Figure 3.**
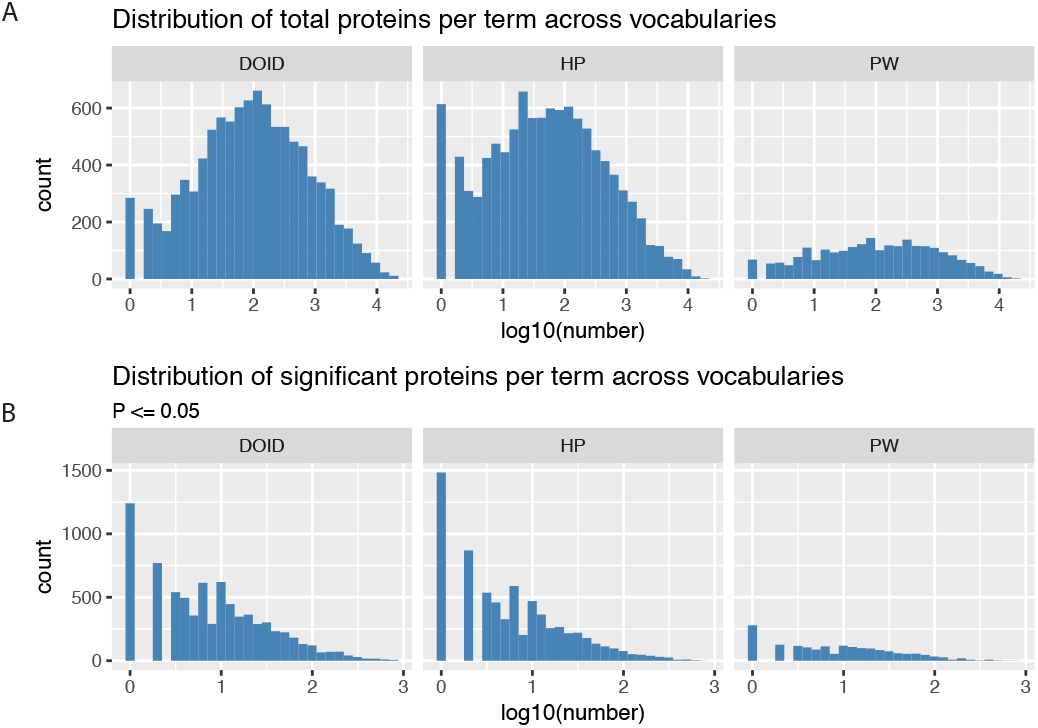
Distribution of total proteins per term in three vocabularies. **A** Number of proteins per term across three queried vocabularies. **B** The distribution of number of significantly associated proteins per term in each vocabulary (*P* ≤ 0.05).

### Reverse query from proteins to significantly-associated topics

Having compiled the prioritized protein lists compiled for a virtually complete collection of human diseases and phenotypes, we explored whether reverse queries could be made from proteins to retrieve information on disease vocabulary terms, i.e., given a protein name, return all the disease areas in which this protein is intensively studied based on literature records. For example, one of the most highly investigated proteins in the topic heart is troponin I (TNNI3). Reverse query with TNNI3 against precompiled popular protein lists of DO and HP terms indicate that as expected TNNI3 is also highly associated with a cluster of cardiovascular-related topics, ranging from myocardial infarction (DOID accession 5844; P: 9.6e-5) to hypertrophic cardiomyopathy (DOID accession 11984; P: 4.1e-3). Utilizing the reverse search strategy on the list of popular disease phenotype proteins above, we find that the top fibrosis protein TGFB1 is significantly associated with mesenchymal cell neoplasm (DOID accession 3350; P:0.059), collagen diseases (DOID accession 854, P: 0.0016), as well as a number of fibrotic diseases including pulmonary fibrosis (DOID accession 3770; P: 0.0045), renal fibrosis (DOID accession 50855; P: 0.0042), and liver cirrhosis (DOID accession 5082; P: 0.031), consistent with its involvement in common disease processes. Moreover, we determined with which other disease terms is another top fibrosis protein CTGF also popularly associated and identified a broad spectrum of disease terms including “connective tissue benign neoplasm”, “connective tissue cancer”, “renal fibrosis”, “liver cirrhosis”, and “scleroderma”. In the Human Phenotype Ontology database, TGFB1 is further associated with phenotypes including cirrhosis, beta-cell dysfunction, hepatic, pulmonary, and renal fibrosis. The pathways associated with TGFB1 includes transforming growth factor *β* signaling pathway, cell-extracellular matrix signaling pathway, peptide and protein metabolic process. Reverse interrogation using precompiled NCD values with the Brenda tissue ontology (BTO) terms revealed that TGFB1 is preferentially associated with a number of fibroblast-related publications in the literature, including myofibrolasts, lung fibroblasts, and others.

One utility of the reverse query is that the curated popular protein lists across human diseases allows the popular proteins to be used as an annotation source for protein list functional analysis, e.g., given a list of differentially expressed proteins in a proteomics experiment, find out whether the significantly up/down-regulated proteins are enriched in proteins that are intensively researched in a particular disease or disease phenotypes. We implemented a new module (FABIAN) to perform gene enrichment analysis against precompiled popular protein lists. To evaluate the potential utility of this approach, we retrieved a transcriptomics dataset on cardiac failure, containing 5 replicates each of control vs. failing hearts from a rodent model of transverse aortic constriction with apical myocardial infarction (GSE56348) [Lai et al., 2014]. We performed a hypergeometric test to identify enriched annotation terms among differentially expressed protein (limma [Ritchie et al., 2015] adjusted *P* ≤ 0.01) against Gene Ontology biological process terms and the precompiled DOID disease-gene associations (Figure 4). The results show that reverse popular protein queries provide complementary annotations to GO Process terms, e.g., we find enrichment of differentially regulated genes that are intensively researched in DOID “collagen disease” and “cartilage disease” terms, corresponding to enrichment of GO extracellular matrix organization” term; as well as “mitochondrial disease” term which corresponds to GO “mitochondrial electron transport, NADH to ubiquinone”. Moreover, enrichment analysis against DOID suggests significant involvement of genes highlighted in “atrial fibrillation” which was not apparent among the enriched GO terms (Figure 5).

**Figure 4.**
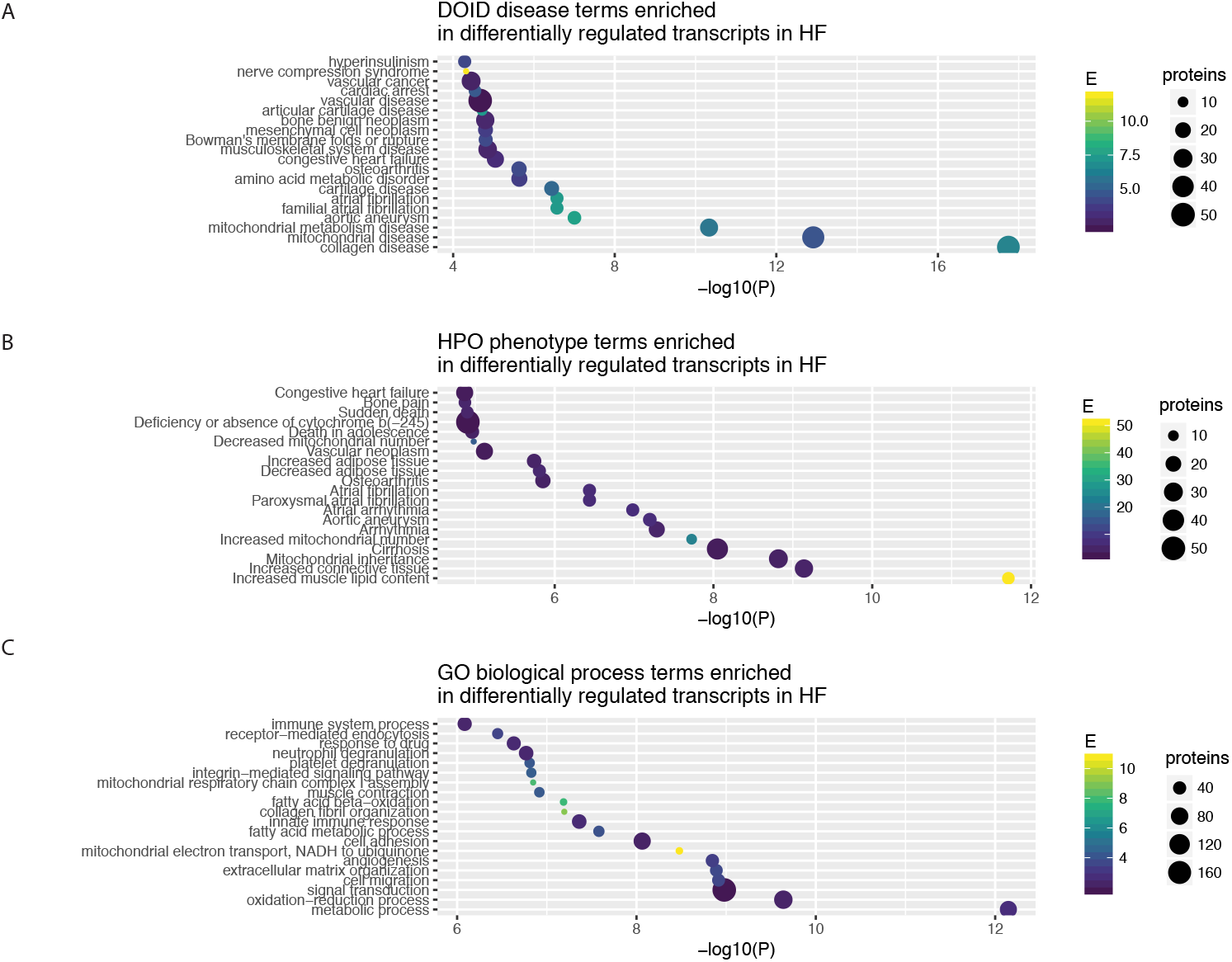
Enriched terms in reverse search (DOID and HPO) vs. Gene Ontology. **A-C** The enriched terms (hypergeometric test *P* ≤ 0.05) from DOID, HPO, and GO associated with differentially expressed genes (*limmaadjusted.P* ≤ 0.01) in a microarray dataset from a rodent model of heart failure.

**Figure 5.**
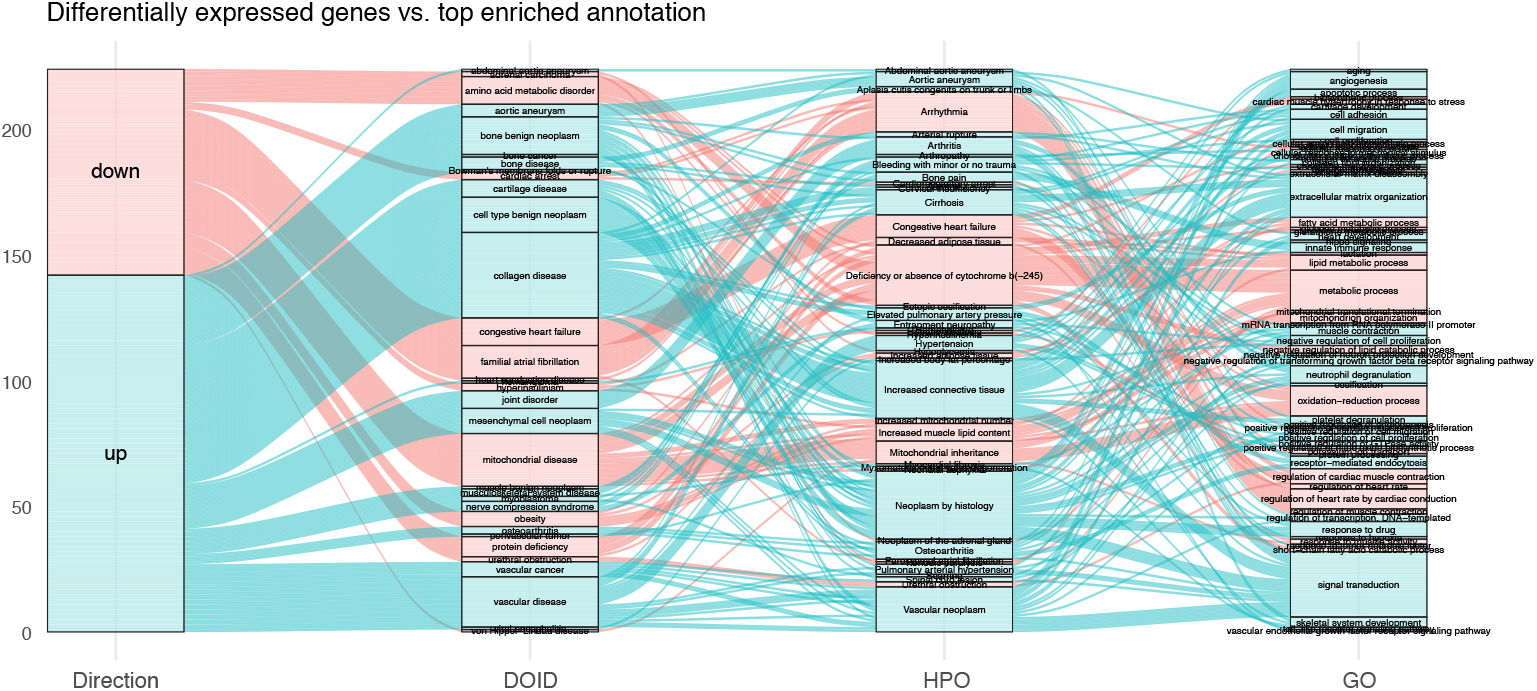
Relationship between assigned DOID, HPO, and GO terms. Top associated terms are shown for each significantly up-regulated (blue) or down-regulated (red) transcripts (limma adjusted *P* ≤ 0.01) in microarray dataset from a rodent model of heart failure. The alluvial streams link the top enriched term of DOID to the corresponding terms in HPO and GO for each transcript. For example, a number of up-regulated transcripts are associated with the “familial atrial fibrillation” term in DOID, corresponding in part to the “arrhythmia” term in HPO, and to the “regulation of heart rate by cardiac conduction” term in GO.

## Discussion

We describe here a method and analysis to prioritize intensively researched proteins associated with cell types, sub-anatomical regions, and molecular phenotypes common across human diseases. Gene/protein prioritization is a recurring informatics problem in biological and biomedical research [Guala and Sonnhammer, 2017]. The gene prioritization problem can be generally stated as follows: given a collection of gene names, identify a subset that is of interest to the topic or disease of interest. For instance, given a list of genes residing at a locus implicated in a genetic mapping study, one may wish to find the causal disease genes or variants responsible for the observed phenotypes. More recently, there has also been interest in protein prioritization efforts to guide rationally allocate limited resources. Such resources may include the prioritized development of research reagents or the distributive fairness in biocuration efforts. The Human Proteome Projects (HPP) within the Human Proteome Organization (HUPO) in particular has the foresight and vision to popularize proteomics assays and reagents, which requires prioritization of genes and proteins that are most worth the effort. This has spawned text-mining and network approaches [Guala and Sonnhammer, 2017, Yin et al., 2017]. Recently we demonstrated that the semantic similarity calculated form the number of publications linked to a gene/protein with a particular topic of interest in the literature may be used to identify priority proteins within a topic within the HUPO HPP. Literature-based gene prioritization is predicated on the hypothesis that over time researchers will choose to work and publish preferentially on proteins relevant to a disease or topic and hence a popular protein will tend also to be biologically important.

Our analysis suggests that regularized co-publication distance metrics offer compatible results to curated disease gene lists, and is distinguished by its compatibility with any topic relevant to PubMed searches. We find that cell types from each organ are preferentially associated with different proteins. The precompilation of popular proteins across disease terms enables a reverse query strategy to identify the diseases and disease phenotypes that have been associated with a protein in the literature. We envision the approach and results presented here will help guide prioritization methods for protein assays, e.g., to develop a panel of MRM assays for fibrosis that can be applicable to ongoing research in the heart, lung, as well as liver. Future work may also leverage the approach discussed here to identify proteins that interact with intensively research proteins but are themselves under-studied in the literature.

## Acknowledgment

This work was supported in part by U.S. Department of Defense grant 16W81XWH-16-1-0592 (J.V.E.), The Barbara Streisand Women’s Heart Center (J.V.E.), The Smidt Heart Institute at Cedars-Sinai Medical Center (J.V.E), The Erika Glazer Endowed Chair in Women’s Heart Health (J.V.E), National Institutes of Health (NIH) research grants P01 HL112730 (J.V.E.), R00 HL127302 (M.P.L.), F32 HL139045 (E.L.), NIH Big Data to Knowledge (BD2K) Program Cloud Credits Model Pilot CCREQ-2017-03-00060 (M.P.L.), and The University of Colorado Consortium for Fibrosis Research and Translation Pilot Grant (M.P.L.).

